# Disruption of the kringle 1 domain of prothrombin leads to late onset mortality in zebrafish

**DOI:** 10.1101/576140

**Authors:** Steven J. Grzegorski, Zhilian Hu, Yang Liu, Xinge Yu, Allison C. Ferguson, Hasam Madarati, Alexander P. Friedmann, Deepak Reyon, Paul Y. Kim, Colin A. Kretz, J. Keith Joung, Jordan A. Shavit

**Affiliations:** Departments of Pediatrics, University of Michigan, Ann Arbor, MI; Departments of Medical Sciences and McMaster University, Hamilton, ON; Departments of Medicine, McMaster University, Hamilton, ON; Departments of Thromosis and Atherosclerosis Research Institute, McMaster University, Hamilton, ON; Departments of Molecular Pathology Unit, Massachusetts General Hospital, Charlestown, MA; Department of Pathology, Harvard Medical School, Boston, MA

## Abstract

The ability to prevent blood loss in response to injury is a critical, evolutionarily conserved function of all vertebrates. Prothrombin (F2) contributes to both primary and secondary hemostasis through the activation of platelets and the conversion of soluble fibrinogen to insoluble fibrin, respectively. Complete prothrombin deficiency has never been observed in humans and is incompatible with life in mice, limiting the ability to understand the entirety of prothrombin’s *in vivo* functions. We have previously demonstrated the ability of zebrafish to tolerate loss of both pro- and anticoagulant factors that are embryonic lethal in mammals, making them an ideal model for the study of prothrombin deficiency. Using genome editing with TALENs, we have generated a null allele in zebrafish *f2*. Homozygous mutant embryos develop normally into early adulthood, but demonstrate eventual complete mortality with the majority of fish succumbing to internal hemorrhage by 2 months of age. We show that despite the extended survival, the mutants are unable to form occlusive thrombi in both the venous and arterial systems as early as 3-5 days of life, and we were able to phenocopy this early hemostatic defect using direct oral anticoagulants. When the equivalent mutation was engineered into the homologous residues of human prothrombin, there were severe reductions in secretion and activation, suggesting a possible role for kringle 1 in thrombin maturation, and the possibility that the F1.2 fragment has a functional role in exerting the procoagulant effects of thrombin. Together, our data demonstrate the conserved function of thrombin in zebrafish, as well as the requirement for kringle 1 for biosynthesis and activation by prothrombinase. Understanding how zebrafish are able to develop normally and survive into early adulthood without prothrombin will provide important insight into its pleiotropic functions as well as the management of patients with bleeding disorders.

**Key Points:**

- Disruption of the kringle 1 domain of prothrombin leads to severe impairment of biosynthesis, activation, and activity
- Prothrombin deficiency is compatible with normal development in zebrafish but leads to inability to form clots followed by early mortality

## Introduction

Maintaining blood flow in a closed circulatory system requires a delicate balance between pro- and anticoagulant factors. Disequilibrium of these factors in either direction can lead to pathology. In response to vascular injury, the balance shifts to procoagulation in an effort to stabilize blood clots and prevent exsanguination. Critical to this clot stabilization is the activation of the zymogen prothrombin, a vitamin K-dependent clotting factor, to form the central clotting serine protease thrombin^1^. Thrombin cleaves soluble fibrinogen into fibrin monomers, which then polymerize to form the insoluble fibrin clot^2^. Human *F2* variants have been linked to both thrombophilia and bleeding diatheses. The most common variant (∼2% in European populations^3^) is a guanine to adenine transition in the 3′ untranslated region at position 20210 that leads to increased plasma prothrombin levels and a 2-3 fold elevated risk of deep vein thrombosis^4,5^. Congenital prothrombin deficiencies are rare; however, acquired deficiencies due to liver failure or vitamin K deficiency are clinically relevant^6,7^.

Structurally, prothrombin consists of six domains: signal peptide, propeptide, Gla domain, kringle 1, kringle 2, and a serine protease domain that consists of a light and heavy chain^1^. Following translation, the propeptide targets the protein for post-translational gamma-carboxylation of the glutamic acid residues within the Gla domain. This vitamin K-dependent process is necessary for proper localization of the mature zymogen to the membrane surface^8^. While relatively understudied, the kringle domains are thought to interact with activated factor V (FVa) during assembly and catalytic turnover of the prothrombinase complex (FVa and activated factor X (FXa))^9^–^11^. Cleavage of prothrombin by the prothrombinase complex results in fragment 1.2 (the Gla and 2 kringle domains) and thrombin (consisting of light and heavy chains). Fragment 1.2 is able to interact with exosite 2 of thrombin to modulate its activity^12^.

Targeted mutagenesis of prothrombin in mice demonstrated 50% mortality by e10.5 with hemorrhage associated with suspected defects in yolk sac vasculature. Mutants surviving to birth succumbed to hemorrhage by postnatal day one^13,14^. Targeting of other common pathway factors, including FV and FX, showed a similar pattern of bimodal lethality^15,16^. These data are suggestive of a secondary role beyond the coagulation cascade during development. Later studies revealed that conditional knockout of prothrombin in adult mice led to mortality due to hemorrhagic events within 5-7 days, although there remained a residual level of prothrombin^17^. Combined with the long half-life of prothrombin (60-72 hours)^18^, it is likely that the adult mice did not achieve complete deficiency prior to lethality.

Due to early lethality of complete genetic knockouts of prothrombin in mice, the nuances of thrombin’s role in *in vivo* hemostasis and development are difficult to study. Zebrafish (*Danio rerio*) has advantages for the investigation of early development because of its high fecundity, external fertilization, and transparent development. Embryonic and adult studies have demonstrated the benefits of zebrafish for the study of hemostatic and other human diseases^19,20^. Conservation of the coagulation cascade in zebrafish is well characterized with a variety of techniques and genetic models^21^–^27^. FX knockout (*f10*^*-/-*^) in zebrafish demonstrated that the model is more resistant to extreme disturbances in hemostasis than mammals^28^. Specifically, mutant zebrafish were capable of surviving several months into adulthood with an absence of this crucial element of coagulation, although they ultimately succumbed due to lethal hemorrhage. Notably, evaluation of their vasculature showed no abnormalities in the embryonic period. In order to determine whether this extended survival was due to low level cascade-independent thrombin generation, we used genome editing TALENs (transcription activator-like effector nucleases) to produce a genetic knockout of the prothrombin gene (*f2*) in zebrafish. We show here that loss of prothrombin in zebrafish does not result in severe developmental defects, but does prevent the formation of clots in response to endothelial injury and leads to early mortality at 2-3 months of age. Furthermore, the mutation generated lends insight into the role of the kringle 1 domain in prothrombin biosynthesis and activation.

## Materials and Methods

### Animal Care

Zebrafish were maintained according to protocols approved by the University of Michigan Animal Care and Use Committee. All wild-type fish were a hybrid line generated by crossing AB and TL fish acquired from the Zebrafish International Resource Center. *Tg(cd41:egfp)* fish were used for tracking of fluorescently labeled thrombocytes^29^. Tris-buffered tricaine methanosulfate (Western Chemical) was used for anesthesia during all live animal experimental manipulation and for euthanasia.

### Sequence Conservation Analysis

Protein sequences (NP_000497.1, XP_001165233.1, XP_003639742.1, NP_776302.1, NP_034298.1, NP_075213.2, NP_989936.1, NP_998555.1, NP_001015797.1) were downloaded from the National Center for Biotechnology Information RefSeq database^30^. Homology predicted propeptides were removed and the prothrombin numbering scheme was used relative to the human prothrombin sequence unless otherwise noted. Zebrafish sequence was modified to include an incompletely annotated exon. Sequences were aligned using MUSCLE^31^ and annotated using the Boxshade server (https://embnet.vital-it.ch/software/BOX_form.html).

### Targeted Mutagenesis using TALENs

The *f2* coding sequence was identified in the zebrafish reference genome (Gene ID: 325881)^32^. TALEN constructs were created using fast ligation-based solid-phase high-throughput (FLASH) assembly targeted to exon 6 of the zebrafish genomic *f2* locus and validated as described^33,34^. mRNA was transcribed from plasmids and injected into single cell embryos. The resulting chimeric founders were raised, outcrossed, and offspring screened for deleterious mutations.

### Genotyping of mutant offspring

The *f2*^Δ14^ allele represents a 14 base pair deletion in exon 6 of zebrafish *f2*. Whole embryos or adult fin biopsies were lysed in buffer (10 mM Tris-Cl, pH 8.0, 2 mM EDTA, 2% Triton X-100, 100 ug/mL proteinase K) for 2 hours at 55°C followed by incubation at 95°C for 5 minutes. One microliter of the resulting solution was used as template for gene specific polymerase chain reaction (Table S1). The products were separated by 3.5% agarose or capillary gel electrophoresis.

### Histochemical Analysis

For *in situ* hybridization, DIG-labeled riboprobes (DIG RNA-labeling kit, Roche) were synthesized using 2 day old wild type embryonic cDNA and gene specific primers with T7 or SP6 overhangs (Table S1). Embryos were fixed overnight at 4°C in 4% paraformaldehyde in phosphate buffered saline prior to dehydration. Permeabilization and staining were performed as described^35^. Stained samples were evaluated by phenotype prior to genotyping.

For hematoxylin and eosin staining, juvenile zebrafish (30-89 days) were fixed overnight in 4% phosphate buffered paraformaldehyde and embedded in paraffin. Sagittal sections (3 µm) were collected every 50 µm and stained.

### Single molecule real-time sequencing of RNA

Three larvae each of wild-type, heterozygous, and homozygous mutant genotypes were homogenized in lysis buffer using a 21-gauge syringe. Total RNA was purified from the lysate using the PureLink RNA Mini Kit (Life Technologies) followed by DNAse I treatment (Invitrogen) and cDNA synthesis using the Superscript III First Strand cDNA kit (Invitrogen). Barcoded primers (Table S1) were used to amplify a 918 base pair (bp) region surrounding the predicted deletion. This region included several non-deleterious single nucleotide polymorphisms (SNPs) in downstream exons known to be allelic with the wild-type and mutant alleles. Purified products from the 9 samples were pooled and sent to the University of Michigan Sequencing Core for library preparation and single molecule real-time high throughput sequencing (SMRT, Pacific Biosciences). Circular consensus reads with at least 5x coverage were filtered for full-length single inserts. The resulting 17,061 reads were subsequently sorted by barcode, allelic SNPs, and splice variation.

### Quantitative Real-Time PCR

Total RNA from whole 3 day old larvae was extracted using the RNeasy mini kit (Qiagen) per manufacturer’s protocol. Three pools of twenty-five whole larvae per biological replicate were used per genotype. The quality and concentration of RNA were assessed using a Nanodrop Lite spectrophotometer (Life Technologies, Grand Island, NY). RNA was transcribed to first-strand cDNA using oligo(dT)_12-18_ primer and Superscript II (Invitrogen). The resulting cDNA was used as template for qPCR (Bio-Rad, iCycler) using SYBR Green PCR Master Mix (Applied Biosystems). The expression level of *f2* was normalized to the *18s* gene and significance analyzed using the double delta Ct method as described^36^.

### Laser-induced endothelial injury

Anesthetized zebrafish larvae were embedded in low melting point agarose at a final concentration of 0.8% and oriented in the sagittal position on a glass coverslip. Larvae were then positioned under an Olympus IX73 inverted microscope with an attached pulsed nitrogen dye laser (Micropoint, Andor Technology). For venous injury, ninety-nine laser pulses were administered to the luminal surface of the endothelium on the ventral side of the posterior cardinal vein (PCV) 5 somites posterior to the anal pore of 3 day post fertilization (dpf) larvae. For arterial injury, pulses were administered to the endothelial surface of the dorsal aorta 3 somites posterior to the anal pore of 5 dpf larvae. Following injury, time to occlusion and/or time to thrombocyte attachment were monitored for 2 minutes. Larvae were manually extracted from agarose for genotyping.

### Chemical treatment

Dabigatran etixalate, apixaban, and rivaroxaban were dissolved in DMSO and diluted in embryo water to final concentrations of 50, 100, and 250 µM, respectively. At 5 dpf larvae were treated for 24 hours prior to laser-mediated endothelial injury on day 6. For ε-aminocaproic acid (Sigma) treatment, zebrafish embryos were incubated in working solution (100 mM) at 1 dpf and the time to occlusion was evaluated at 3 dpf following the laser-induced endothelial injury as previously described^28^.

### o-dianisidine staining

Anesthetized larvae were incubated in the dark for 30 minutes in o-dianisidine staining solution as previously described^37,38^. Larvae were subsequently fixed overnight at 4°C in 4% paraformaldehyde, and pigment was removed in bleaching solution (1% KOH, 3% H_2_O_2_).

### Human prothrombin expression vector construction and injection

Plasmid pF2-bio-His was obtained from Addgene (plasmid #52179) and human *F2* cDNA was amplified using primers containing vector sequence homology (Table S1). pcDNA3.1 was digested with BstBI and KpnI, followed by gel purification of the linearized backbone. The NEBuilder kit was used to fuse the *F2* cDNA with linearized vector and transformed into 10beta competent cells (NEB). Site directed mutagenesis was used for missense mutation introduction^39^ and NEBuilder end homology joining was used to create internal deletions. Replacement of residues 100 to 115 of the human *F2* expression vector with a glutamic acid resulted in a homologous 15 amino acid deletion in prothrombin (Δ15). As the deletion includes the cysteine residue at 114, the effect of introducing a free-cysteine at position 138 due to the loss of its binding partner was investigated by generating two additional vectors that included an alanine point mutation without (C138A) or with the 15-residue deletion (C138A/Δ15). For *in vivo* rescue assays, plasmid DNA (100 pg) in 0.1M KCl was injected into one-cell stage embryos generated from heterozygous *f2*^Δ14^ incrosses using pulled glass capillary needles. Non-viable embryos were removed at 8 hours post fertilization.

### Human prothrombin protein expression and isolation

The constructed cDNAs were transfected into human embryonic kidney 293T (HEK293T) cells to test for their ability to express prothrombin. HEK293T cells were grown to 70-90% confluency in a 6-well plate and then transfected with 5 µg of the constructed DNA (or 5 µL ddH2O) using Lipofectamine 3000 (ThermoFisher Scientific; L3000008). 3.75 and 7.5 µL of Lipofectamine 3000 reagent and 10 µL P3000 reagents were used for transfection. The cells were incubated for 3 days at 37°C, and their media (2 mL) was then collected. The cells were washed with 2 mL of dPBS buffer and the wash buffer was then aspirated. 0.3 mL of 0.25% Trypsin-EDTA was then added to the cells in each well, incubated at 37°C for 5 minutes, then washed with 0.7 mL dPBS buffer and collected separately. The collected cells were centrifuged at 500g for 5 minutes, and the supernatant was discarded. Pelleted cells were suspended in 50 µL RIPA buffer and lysed on ice for 15 minutes before being diluted to 2 mL with sample diluent buffer (HBS-TBS-T20). Prothrombin levels in the expression media and the cell lysate were then quantified by ELISA using a matched-pair antibody set ELISA kit (Affinity Biologicals Inc.; FII-EIA).

The ELISA plate was coated with 1/100 dilution of capture antibody (FII-EIA-C) in coating buffer (50 mM carbonate) at 22°C for 2 hours. The plate was then washed 3 times with PBS-Tween, and the prothrombin samples, including purified wild type with known concentrations, diluted in HBS-TBS-T20, were added and incubated at 22°C for 1 hour. The plate was then washed 3 times with PBS-Tween and the detecting antibody (FII-EIA-D) with 1/100 dilution in HBS-TBS-T20 was added and incubated at 22°C for 1 hour. The plate was then washed 3 times with PBS-Tween, and 50 µL of TMB-ELISA substrate (Thermo Scientific, 34028) was added and incubated at 22°C for 5-10 minutes. 50 µL of 2M sulfuric acid was then added to each well, and the absorbance of the plate was measured at 450 nm using SpectraMax M3 (Molecular Devices), and analyzed using SoftMax Pro (version 5.4).

To characterize the functional characteristics of the prothrombin derivative, the plasmid containing the cDNA for prothrombin C138A/Δ15 was transfected and selected in HEK293T cells to generate a stable cell line (generously provided by Dr. Sriram Krishnaswamy, University of Pennsylvania, USA). Once sufficient numbers of cells were obtained, the cells underwent sustained rolling incubation at 37°C with 5% CO_2_. Expression was induced in D-MEM media supplemented with vitamin K_1_, and was collected every 2-3 days^40^. The media were then centrifuged at 2,000g to remove cell debris, and the supernatant was filtered using a 0.22 µm sterile media filtering system (Millipore). The media was then loaded onto tandem XAD_2_ and Q-Sepharose fast flow columns pre-equilibrated with 0.02M Tris, 0.15M NaCl, pH 7.4 (TBS) and eluted with 0.02M Tris, 0.5M NaCl, pH 7.4 as described previously^41^.

### Prothrombin activation and activity

Prothrombin activation was quantified as described previously^41^. Briefly, Prothrombin (0.1 µM) was incubated with PCPS vesicles (50 µM; 75% phosphatidylcholine, 25% phosphatidylserine), FVa (20 nM), CaCl_2_ (5 mM), and dansylarginine *N*-(3-ethyl-1,5-pentanediyl)amide (1 µM, DAPA) in 0.02 M Hepes, 0.15 M NaCl, pH 7.4 with 0.01% Tween80 (HBST). The reactions were initiated by the addition of FXa (70 pM) and activation was monitored for fluorescence change at 1 minute intervals using a SpectraMax M2 plate reader (Molecular Devices, Sunnyvale, CA). Excitation and emission spectra were 280 nm and 540 nm, respectively, with an emission cutoff filter set at 530 nm. The quantum yield of the thrombin-DAPA complex was determined by plotting the total signal change observed with respect to known concentrations of the thrombin-DAPA complex as described previously^11^.

To determine if the activity of the resulting thrombin from activation of either wild-type or C138A/Δ15 prothrombin were different, two approaches were considered. One approach was to investigate the amidolytic activity of the two thrombin variants using a thrombin-specific chromogenic substrate (S-2238, DiaPharma, West Chester, OH), while the other approach was to test the procoagulant activity of the resulting thrombin in a purified system. They were monitored simultaneously from parallel experiments. Prothrombin was activated as described above by prothrombinase. After 30 min, all prothrombin was activated as verified by SDS-PAGE (not shown). The reactions were initiated by the addition of thrombin (5 nM) to either S-2238 (400 µM) or fibrinogen (3 µM), all in the presence of 5 mM CaCl_2_ in HBST. They were then monitored using SpectraMax M2 at 405 nm, which measures either the cleavage of S-2238 or fibrin clot formation by turbidity change.

### Statistical analysis

The occlusion data were analyzed using Mann-Whitney *U* or two-tailed Student *t* tests. Survival rate was evaluated by log-rank (Mantel-Cox) testing. Significance testing, graphs, and survival curves were made using GraphPad Prism (Graphpad Software, La Jolla, California). P-values (p<0.05 or p<0.0001) were used to evaluate statistical significance.

## Results

### Zebrafish prothrombin demonstrates a high degree of sequence conservation

Following removal of the propeptide, zebrafish prothrombin is a 579 amino acid protein that shares 55% identity and 72% homology with human prothrombin, including 68% identity and 86% homology when comparing the regions corresponding to activated thrombin. Zebrafish prothrombin is detected within the first day of development and leads to measurable fibrin forming activity by 48 hours^42^. The overall structure of prothrombin is highly conserved across species with the presence and spacing of all cysteines and disulfide bonds being preserved (Figure 1).

**Figure 1.**
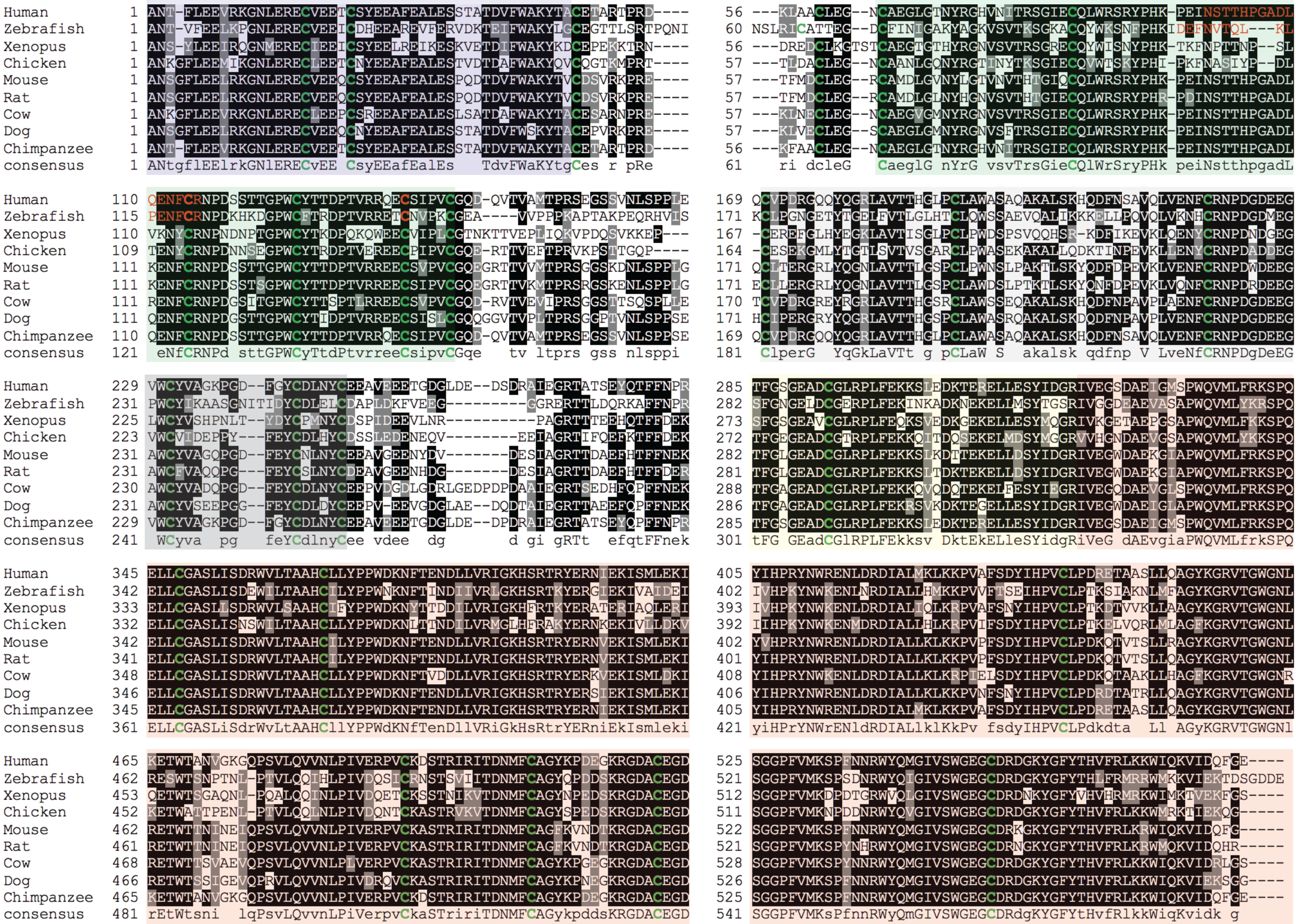
Peptide sequence alignment shows strong conservation of prothrombin across a broad range of species. Green colored residues indicate conserved cysteines. Red colored residues represent amino acids altered by mutagenesis (Δ15; C138A). The prothrombin domains are shaded as follows: Gla (blue), kringle 1 (green), kringle 2 (gray), light chain (yellow), heavy chain (red).

### Targeted genome editing induces a deletion resulting in decreased f2 expression

Utilizing TALEN-mediated genome editing, exon 6 of *f2* was targeted with the aim of creating a frameshift and subsequent nonsense mutation. Sequencing data showed a 14 bp deletion within the region homologous to the human prothrombin kringle 1 domain (Figure 2A). *In situ* hybridization demonstrated decreased, but not absent *f2* expression in homozygous mutants at 3 and 5 dpf compared to wild-type siblings (Figure 2B). This is further supported by qRT-PCR data demonstrating a 60% reduction in transcript in homozygous mutants (Figure 2C). To determine why there was only a partial reduction in expression, we performed deep SMRT sequencing of *f2* cDNA from pooled wild-type and mutant larvae at 3 dpf. Overall, 17,061 consensus sequences were analyzed and 9,097 (61.48%) contained the wild-type allele, all showing the expected splicing pattern. Surprisingly, of the 7,964 (38.52% of total reads) reads from the Δ14 allele, only 53 (0.67%, 0.36% of total reads) contained the 14 bp deletion. The remaining 7,911 (99.33%, 38.16% of total reads) of mutant allele reads did not include the 14 bp deletion, and instead had a 45 bp in frame deletion (Figure 3). Further analysis revealed that there is a cryptic splice acceptor 3′ to the 14 bp genomic deletion in exon 6, resulting in splicing to the exon 5 donor site and a 45 bp deletion in kringle 1 of prothrombin. Notably, this leads to a deletion of 15 amino acids containing a highly conserved cysteine (C119 in the zebrafish peptide) known to form a disulfide bond with zebrafish C143 (Figure 1).

**Figure 2.**
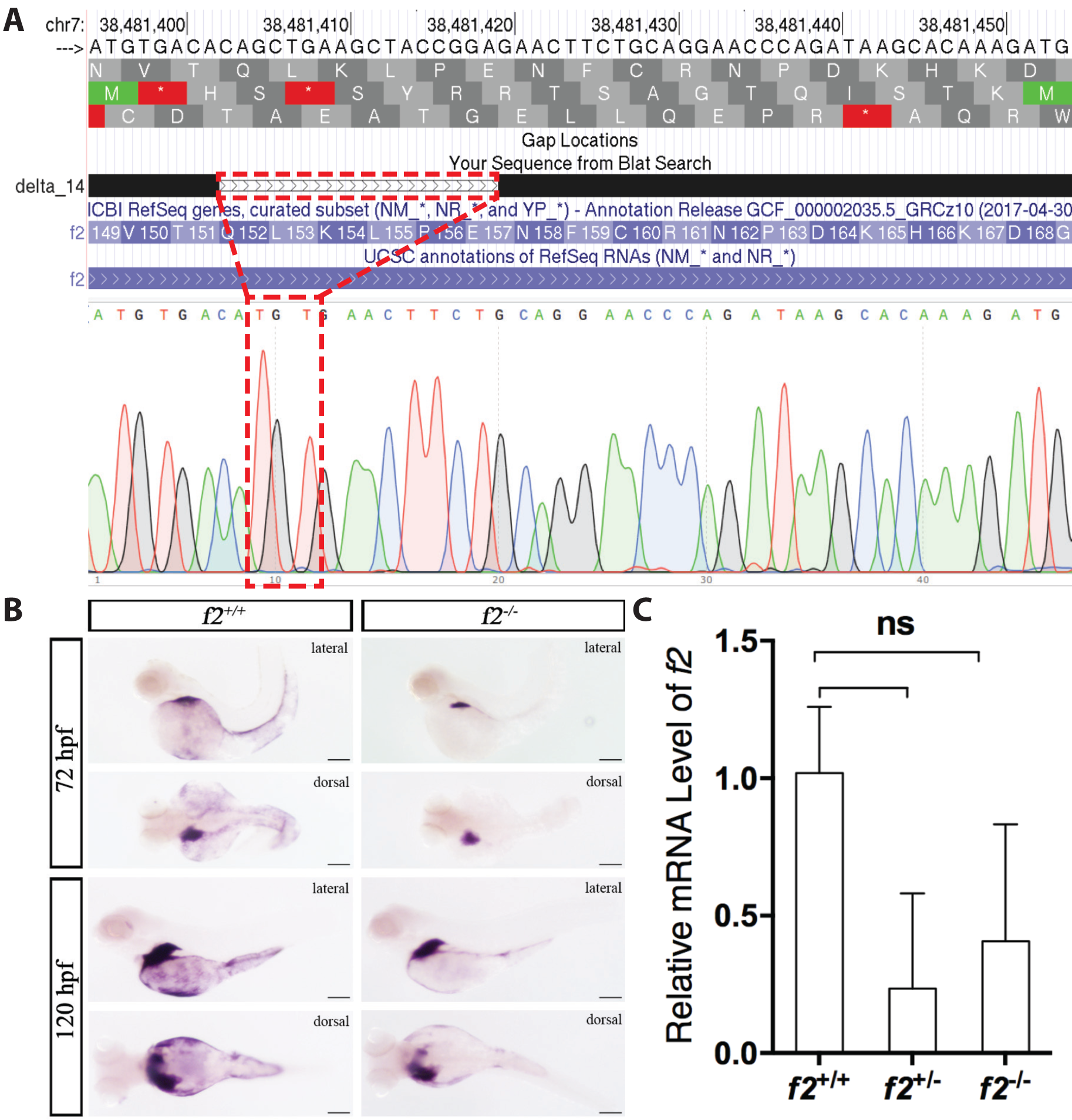
Genome editing creates a 14 bp genomic deletion with a resulting decrease in mRNA expression. (A) Alignment of Sanger sequencing with the chromosome 7 genomic region shows an overall 17 bp genomic deletion replaced with a 3 bp insertion; outlined in red. (B) qPCR data of *f2* expression reveals a non-significant decrease of greater than 50% in heterozygous and homozygous mutants. (C) *in situ* hybridization demonstrates qualitative reduction of transcript at 72 and 120 hours post fertilization in homozygous mutants compared to control siblings. Spatial regulation remains intact with expression restricted primarily to the liver.

**Figure 3.**
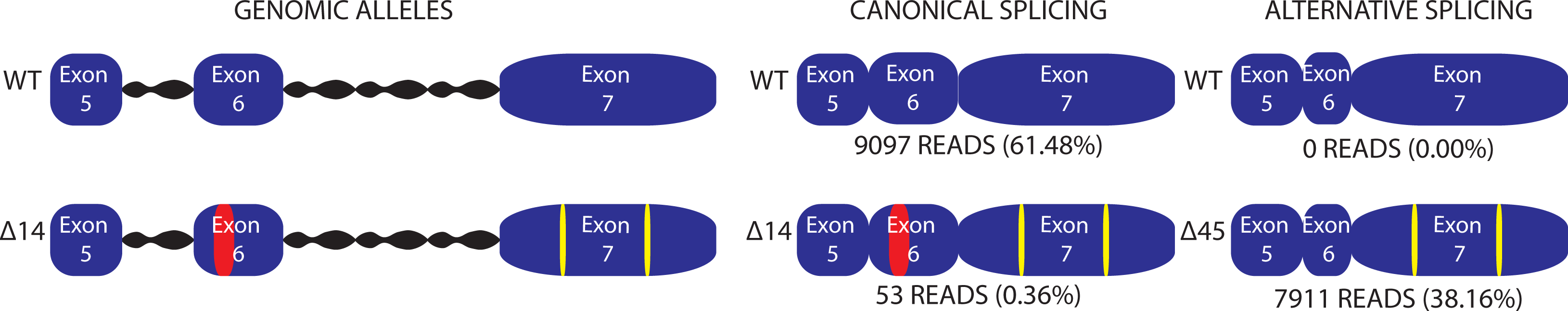
Single molecule real time sequencing of pooled larval mRNA uncovers a cryptic splice site in *f2* following a deletion in exon 6. Allelic SNPs in exon 7 demonstrate that mutant transcripts are preferentially spliced to an acceptor after the mutation, resulting in a 45 base pair deletion.

### Disruption of the prothrombin kringle 1 domain results in adult lethality due to internal hemorrhage

To examine the functional consequence of kringle 1 disruption, we evaluated homozygous mutant fish at various stages of development. Beginning around 1 month of age, *f2*^*-/-*^ fish showed signs of overt internal hemorrhage and the majority died by 2-3 months of age (Figure 4A-B). Gross intracranial and fin hemorrhage was visible, and histological analysis demonstrated occurrence of bleeds within the head, jaw, muscle, fins, and around the heart and abdomen (Figure 4C).

**Figure 4.**
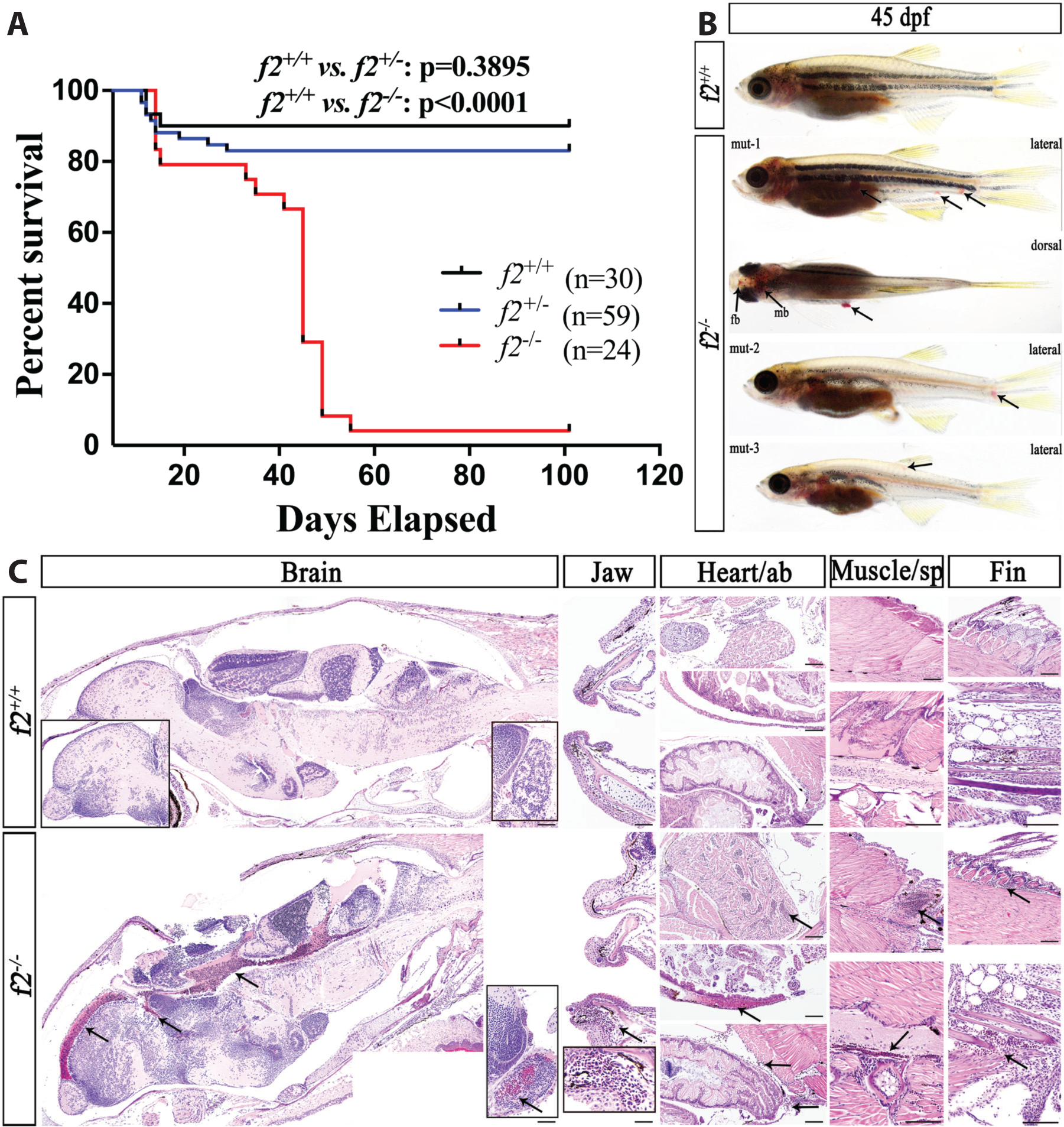
Loss of prothrombin results in early lethality due to hemorrhage. (A) Survival curve demonstrating significant mortality by 2 months of age in *f2*^*-/-*^ siblings. (B) Examples of grossly visible intracranial, intramuscular, and fin bleeds (arrows). fb, forebrain; mb, midbrain. (C) Histological sections of wild-type and *f2*^*-/-*^ siblings demonstrate microscopic bleeds in the brain, jaw, heart, muscle, and fins (arrows).

### Homozygous mutant larvae fail to form thrombi in response to endothelial injury

Given the surprisingly extended survival of homozygous mutants, we checked to see if there was residual detectable thrombin activity in early embryos and larvae. Thrombin has roles in both primary and secondary hemostasis, platelet activation and fibrin formation, respectively. These roles were evaluated using laser-mediated endothelial injury models of arterial and venous thrombus formation. Homozygous mutant larvae were unable to form occlusive thrombi in the venous system in response to injury (Figure 5B). This was refractory to treatment with the fibrinolytic inhibitor ε-aminocaproic acid, further confirming a lack of fibrin generation.

**Figure 5.**
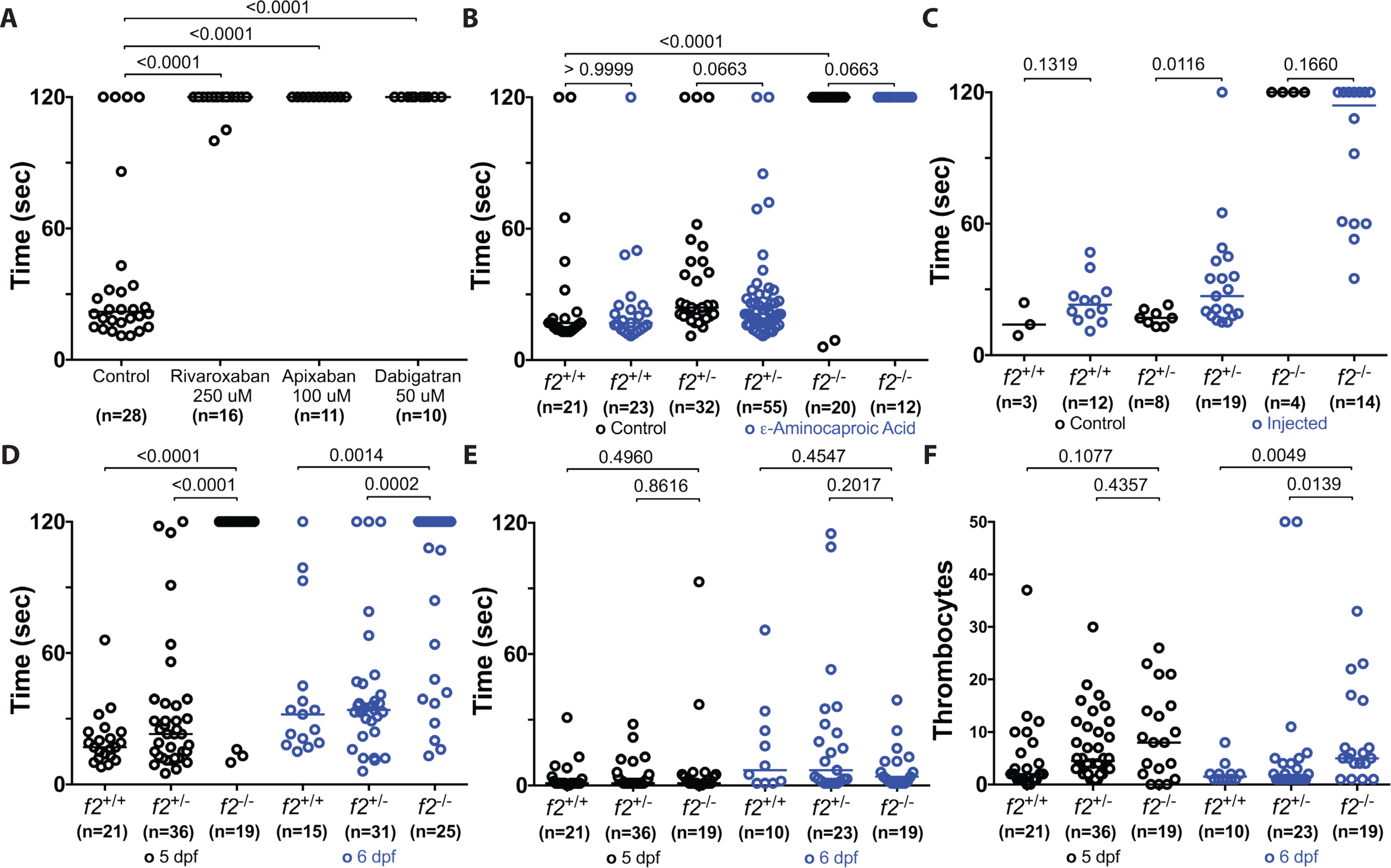
Altered thrombus formation in response to loss of thrombin activity. Larvae were immobilized in agarose, subjected to laser-mediated endothelial injury of the venous or arterial circulation, and followed for 2 minutes by a blinded observer. (A) 24 hour exposure of chemical inhibitors of thrombin (dabigatran) and FXa (apixaban, rivaroxaban) leads to the inability to form occlusive thrombi at 6 dpf in wild-type larvae. Genetic ablation of *f2* results in the inability to form induced PCV thrombi at 3 dpf and is not influenced by inhibiting fibrinolysis (L-aminocaproic acid treatment, blue). Overexpression of human prothrombin cDNA (blue) rescues the ability to form thrombi in the PCV at 3 dpf. (D) Homozygous mutant larvae demonstrate a significant impairment in arterial thrombus formation at 5 and 6 dpf without any changes to the time to initial thrombocyte attachment (E). (F) The number of thrombocytes attached to the site of injury in 2 minutes is significantly increased at 6 dpf in *f2* homozygous mutants.

To validate the specificity of the coagulation defect, embryos generated from a heterozygous incross were injected at the one-cell stage with plasmid expressing human prothrombin cDNA. Endothelial injury at 3 dpf induced clot formation in 50% of homozygous embryos in contrast to uninjected controls (Figure 5C).

Assessment at 5 and 6 dpf in the *Tg(cd41-egfp)* thrombocyte-labeled background showed that larvae have a decreased ability to form occlusive arterial thrombi as well (Figure 2D). The time to thrombocyte attachment was not statistically different between groups while the number of attached thrombocytes appears to be increased in the *f2*^-/-^ larvae (Figure 2E-F). Overall these data reveal a deficiency in primary and a loss of secondary hemostasis. Despite this, o-dianisidine staining revealed no overt signs of hemorrhage in 1 week old wild-type or homozygous mutant larvae (data not shown).

### Zebrafish thrombin activity is inhibited by direct oral anticoagulants

Reduction of vitamin K epoxide to vitamin K is necessary for the carboxylation and subsequent activity of many coagulation factors, and the inhibition of this process by warfarin has been validated in zebrafish^24,43,44^. To investigate how anticoagulation using the new direct oral anticoagulants affects zebrafish hemostasis, three agents were tested: direct thrombin inhibitor (dabigatran) or direct FXa inhibitors (rivaroxaban and apixaban). Following 24 hours of treatment, 6 dpf larvae exposed to dabigatran etexilate, rivaroxaban, and apixaban were unable to form occlusive thrombi in response to injury (Figure 5A). These data show that the carboxylesterase function required to convert dabigatran etexilate to dabigatran is conserved in zebrafish, as is the ability to inhibit zebrafish thrombin. Apixaban and rivaroxaban also demonstrate conserved inhibition of FXa, which consequently inhibits prothrombin activation. Importantly, no gross hemorrhage was observed, confirming that zebrafish are able to tolerate loss of thrombin and FXa activity in the embryonic/larval period without bleeding, unlike the mouse knockouts.

### An intact kringle 1 domain is necessary for proper secretion and procoagulant activity

To interrogate how the loss of the kringle 1 domain alters prothrombin function, three orthologous variants of human prothrombin (Δ15, C138A, and C138A/Δ15) were investigated. As the deletion includes C114, the effect of introducing a free-cysteine at position 138 due to the loss of its binding partner was investigated by generating two additional vectors that include an alanine point mutation without (C138A) or with the 15-residue deletion (C138A/Δ15). Following transient expression in HEK293T cells, wild-type prothrombin was secreted at levels greater than 11 ng/mL (Figure 6A). In contrast, all 3 mutant plasmids secreted less than 2 ng/mL and were not significantly different than negative controls. These data suggest that the loss of these residues within the kringle 1 domain adversely impact protein folding and/or secretion.

**Figure 6.**
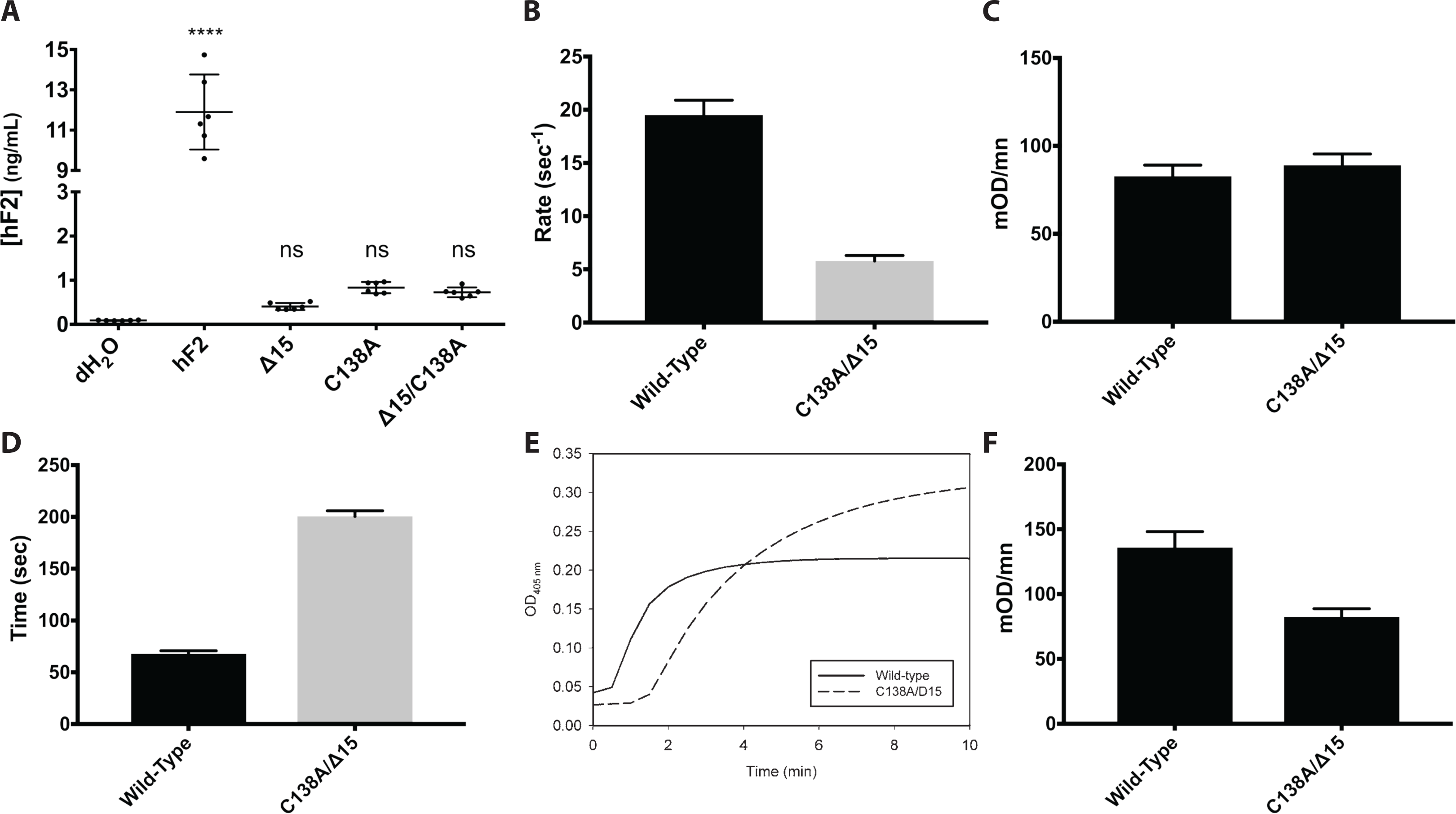
Expression and activation of prothrombin variants, and the resulting activity of thrombin upon activation. (A) Expression of prothrombin variants in HEK293T cells results in reduced secretion levels that are indistinguishable from control. (B) Rates of prothrombin activation by prothrombinase measured using DAPA. Once prothrombin variants were fully activated by prothrombinase, their ability to cleave (C) synthetic substrate S-2238, or (D) fibrinogen were monitored. (E) Clotting profile between wild-type and C138A/Δ15 demonstrate (i) clot time differences are largely due to delayed clot initiation, (ii) total turbidity change differs, suggesting altered thrombin activity. (F) The rates of clot formation were determined from the clotting profile.

To assess whether there is a sustained low level of prothrombin secretion with similar activation kinetics that may afford the extended mutant survival, the prothrombin derivative C138A/Δ15 was recombinantly expressed and isolated. The quantum yield of fluorescence signal change observed upon thrombin-DAPA complex formation was comparable between wild-type (3262 ± 347 RFU/µM) and C138A/Δ15 (3354 ± 180 RFU/µM), consistent with the protease domain remaining intact. When the initial rates of activation by prothrombinase were quantified using DAPA, wild-type prothrombin was 3.3-fold faster (19.5 ± 1.4 s^-1^) compared with C138A/Δ15 (5.8 ± 0.5 s^-1^) (Figure 6B).

The functionality of thrombin generated from the two prothrombin variants was then investigated. Consistent with the similar quantum fluorescence yield values observed from the DAPA-thrombin complex, the amidolytic activity of thrombin towards S-2238 chromogenic substrate was similar between the wild-type (82.7 ± 6.5 mOD/min) and C138A/Δ15 (89.0 ± 6.4 mOD/min) (Fig. 6C). Again, this is consistent with the fact that the mutations are confined to the kringle 1 domain. When the ability to cleave fibrinogen was investigated, however, C138A/Δ15-thrombin had a significantly increased clot time (200.6 ± 5.3 s), mostly due to delayed clot initiation (Figure 6D), as well as a 1.5-fold higher total turbidity change (Figure 6E) compared with wild-type (67.7 ± 3.1 s). The rate of clot formation was also slower (82.4 ± 6.3 mOD/min) compared with the wild-type (135.9 ± 12.3 mOD/min) (Figure 6F). These data suggest that thrombin from C138A/Δ15 prothrombin has a lower thrombin-like activity towards exerting its procoagulant function, which is also consistent with the delayed clot initiation coupled with higher total turbidity change.

## Discussion

The zebrafish model has been developed as a useful tool for understanding coagulation, especially during early development. In contrast to mammals, in which a number of coagulation factors are necessary for embryonic and/or neonatal viability, zebrafish are able to survive the loss of many aspects of the canonical cascade at least until early adulthood. Loss of antithrombin, fibrinogen, FV, and FX are all compatible with development to adulthood^22,26^–^28,45^. Targeting of *f10* in fish results in the absence of larval hemostasis, but is accompanied by extended survival^28^. These data suggested that there could be residual thrombin activity present. Previous studies have utilized transient knockdown and chemical inhibition to study the loss of prothrombin in zebrafish^42,44,46,47^. These studies provide valuable insight into the conserved function of prothrombin and suggest a potential role for the coagulation cascade in development. Unfortunately, these technologies are susceptible to toxicity and off target effects. Additionally, specific chemical inhibition targets the proteolytic activity of thrombin without addressing the potential for exosite mediated binding of other factors. We sought to leverage knockout technology to create a specific genetic knockout of *f2*. We produced a genomic deletion in exon 6 with the intent of creating a nonsense mutation. Unexpectedly, we found reduced expression of an alternatively spliced version of prothrombin lacking a conserved cysteine in the kringle 1 domain. *In vitro* biochemical studies demonstrate that this protein is synthesized and secreted at a negligible level, and any secreted protein has decreased activation and activity. *In vivo*, the Δ15 allele results in the loss of the ability to form fibrin rich clots and lethality by 2 months of age due to spontaneous hemorrhage. However, it appears that thrombocyte attachment is unaffected, consistent with the dogma that platelet adherence is mediated primarily via von Willebrand factor^48^–^50^. Overall, our data suggest that this is a null allele, although we cannot completely rule out residual thrombin activity. However, the results are very similar to the *f10* knockout, suggesting that both represent complete loss of thrombin.

Surprisingly, no obvious developmental defects were observed, including grossly normal vascular development, with survival through early adulthood. Previous work using antisense technology has demonstrated a variety of developmental malformations including hemorrhage, and circulatory, brain, and tailbud malformations^46^. These defects were not observed across multiple clutches in our knockout. This inconsistency could be due to off target effects of antisense knockdown or suggest the possibility of a role for maternally-derived prothrombin. However, this could also be due to potential genetic compensation that can occur in response to genomic editing^51^. Notably, the *victoria* mutant, which mapped close to *f2* and is likely a mutant allele, similarly affects induced thrombus formation without a described developmental defect^52^, consistent with our data.

Kringle 1 is an understudied domain of prothrombin, but it is thought to directly bind to FVa and inhibit FXa catalyzed prothrombin activation^10^. Our data have shed new light on the role of this domain in thrombin activation and function. The impairment of thrombin activity occurs at multiple levels. First, presumed inefficiency of the cryptic splice site in mRNA maturation leads to a roughly 60% reduction in transcript levels. Second, when introduced into human prothrombin, the deletion resulted in a substantial decrease in secretion. This could be explained simply by misfolding, or may be indicative of a more involved role of the kringle 1 domain in regulating prothrombin secretion as has been previously suggested^53^. Finally, we looked for any residual thrombin activity by examining activation and activity of the mutant. C138A/Δ15 prothrombin was shown to have a 3-fold reduction in activation, the thrombin it produced had a 2-fold reduction in its ability to cleave fibrinogen, while maintaining a functional active site as demonstrated by its preserved amidolytic activity towards a synthetic substrate. This is not surprising as the mutation did not impact the protease domain of thrombin. In addition, our data suggest that the F1.2 fragment has a functional role in exerting the procoagulant effects of thrombin in fibrinogen cleavage. Overall, the compounding reductions in transcription, secretion, activation, and activity suggest that this deletion is a complete loss-of-function, while at the same time uncovering a potential role for the kringle 1 domain in the eventual maturation of prothrombin.

In comparison to mammals, the ability of zebrafish to survive the absence of thrombin into adulthood is surprising. However, the zebrafish literature repeatedly demonstrates a resistance to severe coagulopathies (loss of antithrombin, prothrombin, FV, FX) ^22,28,45^ that cause embryonic and/or neonatal lethality in mammals. This may be due to a combination of influences, including the absence of birthing trauma, limited hemostatic challenges in an aqueous laboratory environment, a relatively low systolic blood pressure, and possible species-specific genetic differences. Nevertheless, the phenotypes of all of these mutants eventually converge with their mammalian counterparts in adulthood. Overall, this study further demonstrates the conservation of the coagulation cascade in fish while leveraging unique physiologic differences to build our understanding of the complex biological functions of thrombin. Furthermore, understanding how fish tolerate a such a severe bleeding diathesis could provide new insight into managing patients with congenital bleeding disorders or acquired hemorrhage, with future studies leveraging the model system to develop new diagnostic markers and therapies.

## Acknowledgements

The authors thank the University of Michigan Sequencing and Microscopy & Image Analysis cores for services. This work was supported by National Institutes of Health grants R01-HL124232 and R01-HL125774 (J.A.S.), and T32-GM007863, T32 HL125242, and an American Heart Association Predoctoral Fellowship Award (S.J.G.), a Hemophilia of Georgia Clinical Scientist Development Grant, the National Hemophilia Foundation/Novo Nordisk Career Development Award, and the Bayer Hemophilia Awards Program (J. A. S.), and NIH R01 GM088040 (J.K.J.). J.A.S. is the Diane and Larry Johnson Family Scholar of Pediatrics. This work was also supported by Natural Sciences and Engineering Research Council of Canada (RGPIN-2017-05347 to P.Y.K.). P.Y.K. holds the Department of Medicine Early Career Award (McMaster University).

## Authorship

### Contribution

S.J.G. designed and performed research, analyzed data and wrote the manuscript; Z.H. designed and performed research, analyzed data and assisted with figure design. C.K. designed and performed research, and analyzed data; D.R. and J.K.J designed and performed research; Y.L., X.Y., A.C.F, H.M., and A.F. performed research. P.K. and C.A.K. designed research, analyzed data, and edited the manuscript; J.A.S. designed, performed, and supervised research, analyzed data, and wrote the manuscript.

